# Using machine learning to automate the analysis of an olfactory habituation-dishabituation task in mice

**DOI:** 10.64898/2026.02.24.706573

**Authors:** S. Boyanova, M. H. Correa, R. S. Bains, F. K. Wiseman

## Abstract

**Introduction:** Improving the efficiency and accuracy of annotation and extraction of performance data from mouse behavioural tasks will improve both the throughput and scientific value of preclinical research.

**Methods:** Here, we present and validate an automated pipeline for the annotation and quantification of performance in a mouse olfactory habituation-dishabituation task, using a single side-view camera, resulting in occluded body parts. We created a pipeline for task analysis, combining DeepLabCut, for pose-estimation, and SimBA, for behavioural classification to automatically quantify odour interaction (sniffing time) in a three-odour (water, familiar mouse social odour, novel mouse social odour) variant of the task. We used a subset of previously published, fully manually annotated datasets to train the models and unseen videos from the same study to validate the utility of our machine learning pipeline.

**Results and conclusion:** Our analysis pipeline estimated behavioural performance in the task with high accuracy, and the data produces similar technical and biological results to manual methods when analysed by linear mixed modelling. Thus, we validated the utility of our new pipeline for the automated scoring of this mouse sensory task.

## Introduction

Behavioural analysis is fundamental to the assessment of preclinical mouse models of neurological disease (Von Ziegler, Sturman, & Bohacek, 2021). Sensory inputs and motor outputs are critical to the manifestation of behaviour and cognitive processes and can be independently impacted by disease (Ison, Allen, & O’Neill, 2007; Rodgers, Born, Das, & Jankowsky, 2012; Ryan, Young, Moy, & Crawley, 2008). Thus, battery assessment of both clinically relevant behaviours/cognitive domains and associated sensory and motor processes is important for the assessment of new lines, and determining the potential translational impact of interventions. Olfaction is a key sensory input in mice and is particularly important for the interpretation of social tasks, including the Crawly three chamber test of social motivation and memory (Ryan et al., 2008). Olfaction can be quantified in mice using the habituation-dishabituation task, to test the animal’s ability to differentiate between different odours; by measuring the amount of time an animal interacts with (sniffs) repeated presentation of a series of odours. Novel odours typically elicit increased interaction, which wanes (habituates) over time.

Recent machine learning (ML) methods to automate the quantification of behaviour permit a dramatic increase in analytical throughput and accurate measurement of a greater range of behaviours. A wide variety of ML tools capable of body parts pose estimation, such as DeepLabCut, (Mathis et al., 2018; Pereira et al., 2022), and behavioural classification (Chan et al., 2025; Marks et al., 2022), such as SimBA (Goodwin et al., 2024) have been developed. DeepLabCut is a pose-estimation framework based on deep convolutional networks commonly including ResNets, pretrained on the large-scale object recognition benchmark ImageNet. The ResNet outputs spatial probability densities, the probability density of each body part represents the ‘evidence’ that this body part is at a particular location (Mathis et al., 2018). To then generate meaningful behavioural quantification, the outputs from DeepLabCut, are further processed using a ML behavioural classifier to determine the duration or frequency of behaviours of interest. The ML tool used to train behavioural classifiers in this study, SimBA, uses random forest machine learning classifiers for behavioural predictions by computing explainable feature representations of movement, angles, paths, velocities, distances and sizes within individual frames and as rolling time window aggregates (Goodwin et al., 2024).

The most frequent use case of such ML tools for mouse behavioural analysis focuses on, top-view tracking in conventional tasks. However, in some tasks, much of the complexity of animal behaviours is better captured using a side-view projection, for example, the interaction of an animal with odour cues in the olfaction habituation-dishabituation task. However, the use of a single side-view camera creates a technical challenge caused by the occlusion of body parts, and difficulties with depth estimation. Furthermore, most mouse behavioural data is recorded in black and white using an infra-red or near infra-red camera. Therefore, here we selected two of the most mature and validated ML tools in the behavioural community - DeepLabCut (Mathis et al., 2018) and SimBA (Goodwin et al., 2024) to develop an automated scoring pipeline for the olfaction habituation-dishabituation task. We present and validate this new pipeline for the automated quantification of odour cue interaction (sniffing) using side-view greyscale videos of the habituation-dishabituation task in laboratory mouse models of human disease. We used a manually annotated (ground-truth) dataset, of mice undertaking the olfaction habituation-dishabituation task as part of a previously published longitudinal test battery (Boyanova et al., 2025). This study used two independent lines of genetically modified animals that model aspects of ALS/FTD (*C9orf72^GR400/+^*and *Tardbp^Q331K/Q331K^*). This new pipeline can be used or adapted to automate the quantification of the olfaction habituation-dishabituation task using side-view projection of odour cue presentation, facilitating rapid and accurate data production for the study of other animal models of disease and fundamental biology.

## Methods

Primary data for this paper was previously published in Disease Models and Mechanisms (Boyanova et al., 2025), methods for data acquisition are reported below.

### Animal welfare and husbandry

All animals were housed and maintained in the Mary Lyon Centre at MRC Harwell under specific pathogen-free (SPF) conditions, in individually ventilated cages (IVC) adhering to environmental conditions as outlined in the Home Office Code of Practice. All animal studies were licensed by the Home Office under the Animals (Scientific Procedures) Act 1986 Amendment Regulations 2012 (SI 4 2012/3039), UK, and additionally approved by the Institutional Ethical Review Committees. Mice were randomised, blocked by genotype and sex at the time of weaning, into cages of 3 to 5 mice. All mice used in the study were bred in the Mary Lyon Centre at MRC Harwell and were housed in individually ventilated cages (IVCs; Tecniplast BlueLine 1284), on grade 4 aspen wood chips (Datesand, UK), with shredded paper shaving nesting material and small cardboard play tunnels for enrichment. The mice were kept under controlled light (light 07:00–19:00; dark 19:00–07:00), temperature (22 °C ± 2 °C) and humidity (55% ± 10%) conditions. They had free access to water (25 p.p.m. chlorine) and were fed *ad libitum* on a commercial diet (SDS Rat and Mouse No.3 Breeding diet (RM3).

### Animal genetics

The generation of the *C9orf72^em2.1Aisa^* (MGI:6827370), here called the *C9orf72^GR400/+^* mouse model is described in Milioto et al 2024 (Milioto et al., 2024). The *C9orf72^GR400/+^* line was maintained on a C57BL/6J background by heterozygote backcross before the generation of phenotyping cohorts. Phenotyping cohorts were generated by crossing either male or female *C9orf72^GR400/+^* heterozygotes to C57BL/6J wildtypes (WT). The generation of the *Tardbp^em1Rhbr^* (MGI:6157626), here called the *Tardbp^Q331K/Q331K^* mouse model is described in White et al 2018 (White et al., 2018). The *Tardbp^Q331K/Q331K^* line was maintained on a C57BL/6J background by heterozygote backcross prior to the generation of phenotyping cohorts. Phenotyping cohorts were generated by heterozygote intercross to generate WT and *Tardbp^Q331K/Q331K^* homozygote animals.

### Genotyping

DNA was extracted from ear biopsy, isolated at postnatal day (P)14 using TaqMan Sample-to-SNP (Applied Biosystems). Mice were genotyped for *C9orf72^GR400^* using TaqMan WT and mutant quantitative PCR assays duplexed with Dot1l reference allele. The following primers and probes were used for the *C9orf72^GR400^* mutant allele (forward, 5′-TTCCAGATTACGCTTACCATAC-3′; reverse, 5′-CGACCTCTTCCTCGTCCT-3′) and probe (5′-FAM-TACCTCGTCCACGTCCTCGTCTTC-BHQ1-3′), *C9orf72^+^* WT allele (forward, 5′-CTATTGCAAGCGTTCGGATAATG-3′; reverse, 5′-CTTGGCAACAGCAGGAGAT-3′) and probe (5′-FAM-TGGAATGCAGTGAGACCTGGGATG-BHQ-3′), and reference Dot1l allele (forward, 5′-TAGTTGGCATCCTTATGCTTCATC-3′; reverse, 5′-GCCCCAGCACGACCATT-3′) and probe (5′-VIC-CCAGCTCTCAAGTCG-MGBNFQ-3′). Mice were genotyped for the *Tardbp^Q331K^* alleles using allelic discrimination assays, using a common pair of primers for both *Tardbp* alleles (forward, 5′-TCTGCTGGCTGGCTAACAT-3′; reverse, 5′-GGGTGGAGGGATGAACTTTG-3′). To discriminate between the mutant and WT alleles, different probes were used (TardbpQ331K, 5′-TET-AACTGCTCTTCAACGCT-BHQ1-3′; *Tardbp^+^*, 5′-FAM-CAACTGCTCTGCAACG-BHQ-3′).

### Olfaction task

The olfaction test was run at 15- and 67-weeks of age. The number of mice in the *C9orf72* study at 15-weeks of age were WT=24 (female = 12, male = 12) *C9orf72^GR400^*^/+=^24 (female = 12, male = 12), at 67-weeks of age WT=20 (female = 11, male = 9), and *C9orf72^GR400^*^/+^=18 (female = 9, male = 9) where two videos were excluded due to wrong food hopper being used for the task due to a technical error. In the *Tardbp* study at 15-weeks of age - WT = 27 (female = 12, males = 15), *Tardbp^Q331K^*^/*Q331K*^ = 24 (female = 9, male = 15) where 3 mice were excluded due to wrong order of odours presentation during the test, treated as procedural failure, at 67-weeks of age - WT = 21 (female = 11, males = 10), *Tardbp^Q331K/Q331K^* = 19 (female = 10, male = 9), where one 1 mouse was excluded due to corrupted video file.

Mice were removed from their home cage and allowed to acclimatise to a clean IVC placed in a home cage analysis rig (Actual Analytics Ltd., Edinburgh, UK) for 30 minutes prior to the start of the test without access to food but access to water. Videos were recorded through the infra-red camera of the Home Cage Analyser system (Actual Analytics Ltd., Edinburgh, UK). The odours used for this test were water (control), familiar mouse, and novel mouse (social odours) presented on sterile cotton swabs through the access for the water bottle of the IVC. The cotton swabs for water were prepared by pipetting 50 µl of deionised water onto the swab. The social odours were prepared by wiping the cotton swab in a zigzag fashion across the bottom of a used cage, either the animal’s home cage or an unfamiliar cage. After 30 minutes the water bottle was removed, and the first cotton swab was presented in the cage through the water bottle access with the food hopper in place. The mouse was allowed to explore the cotton swab freely for 2 minutes, followed by 1-minute inter-trial interval. Each odour type was presented three times in a row. The order of the odour types was counter-balanced and randomised across the mouse cohorts. The time spent sniffing was scored manually using SimpleVideoCoder (Barto, Bird, Hamilton, & Fink, 2017), the scorer was unaware of the mouse genotype. We scored the time the mouse spent sniffing each cotton bud. Sniffing was defined as orientation of the mouse’s head and nose towards the cotton bud, and distance of the nose at least 1 cm away from the front and the bottom of the food hopper, as well as the lower one third of the back of the food hopper. Licking or biting of the cotton bud was not included in the time scored. If the mouse spent less than 10 seconds interacting with the first presentation of a given odour, this run and the consecutive two presentations of the same odour were not used in the analysis, since the mouse failed to sufficiently engage with the stimulus on the first presentation (‘Behavioral Models | Behavioral and Functional Neuroscience Lab | Wu Tsai Neurosciences Institute’, 2023).

### Machine learning pipeline training Video preprocessing

For the training stage, a total of 60 videos were randomly selected, balanced for age and genotype – 15 videos per age per genotype from each study (WT and *C9orf72^GR400/+^*, WT and *Tardbp^Q331K/Q331K^*^)^, from a total of 183 videos (25 fps) from the study, previously analysed using manual annotation (Boyanova et al., 2025). To avoid bias, the experimenter was unaware of genotype during this process. Videos were cropped to the region of interest (area proximal to the metal hopper were the odour-cotton buds are presented) (Figure 1). This reduces the size of the videos and complexity of animal behaviour recorded, speeding up downstream processing.

**Figure 1.**
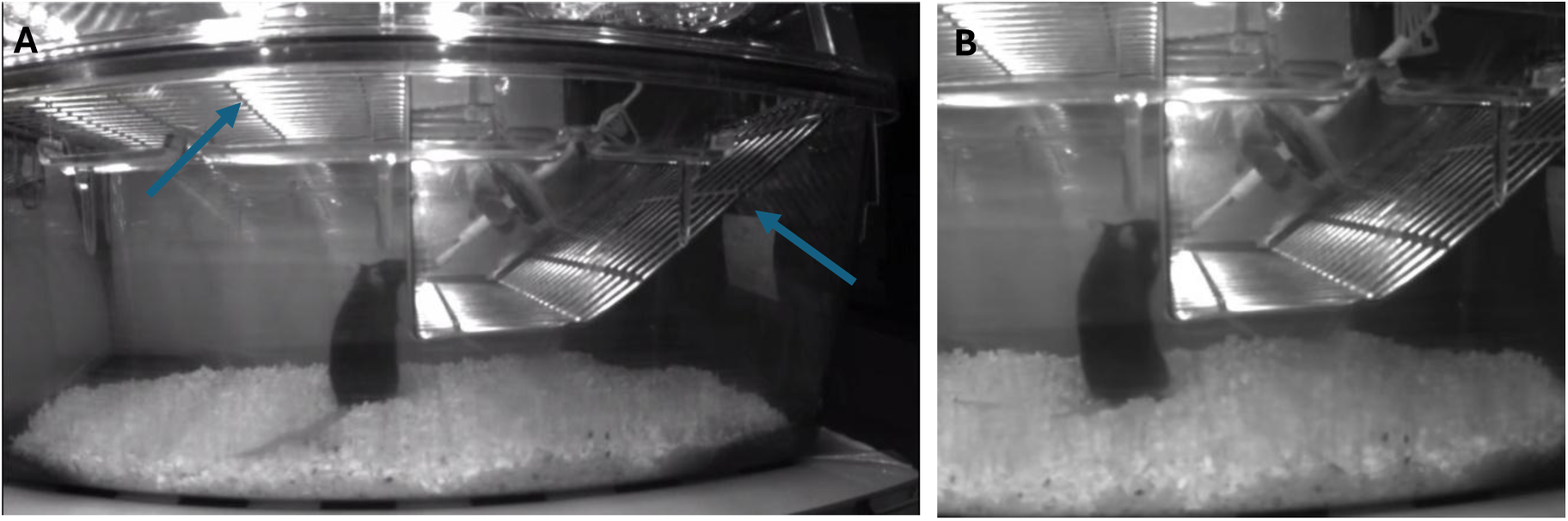
Preprocessing of videos of the habituation-dishabituation task. A) Full frame from an example video and B) Cropped frame used for training and validation on unseen videos in the pipeline. Blue arrows show the hopper bars used as borders to crop the area around the hopper consistently between the videos.

### Pose estimation training using DLC

To track the position of the animal, DeepLabCut (DLC) v. 2.3.9 software (Mathis et al., 2018) was trained using this pre-processed training set. 2069 video frames from the 60 training videos were manually labelled for 10 body parts of interest on the mouse (snout, middle point at eye level, left ear, right ear, back of the neck, front of the neck, two points at the back, base of the tail, and abdomen). To further improve tracking, additional spatial locations relevant to the behaviour of interest were also annotated (four corners of the hopper, the tip of the cotton bud, the checkerboard floor of the arena when visible) (Figure 2 A). To improve training, systematic identification of failed tracking of the animal was implemented. Retraining was focused on these cases, 16 cycles of consecutive addition of extra frames using deep convolutional neural network (CNN) resnet 101, batch size 8, and imgaug image augmentor, was used to improve the outcome. For both training and retraining, the dataset was split randomly (shuffle = 1) into train (95% of frames) and test (5% of frames) subsets, to train and assess network performance, respectively for each cycle. Test error – mean absolute error measured in pixels between the predicted position of the labels and the manual annotation was used to assess network performance. The training cycles were repeated until test error of 2.42 px was reached (spatial resolution of approximately 1.1 mm, p cut-off = 0.6), using 2069 labelled frames, and 2150000 iterations, Nvidia RTX4090 GPU was used during DLC training.

**Figure 2.**
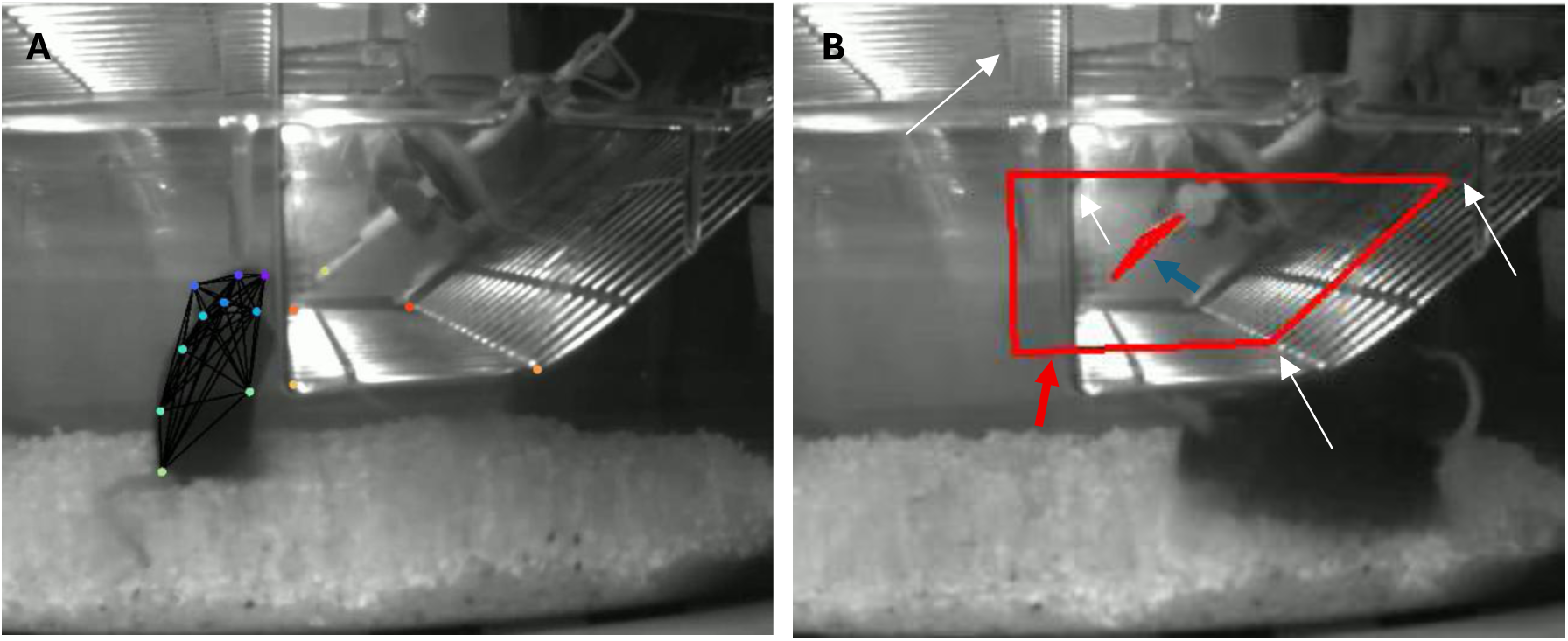
Machine learning annotations for the habituation-dishabituation task analysis pipeline. A) DLC labelling of mouse body parts (green, blue, purple dots), tip of cotton bud presenting odour cue (yellow) and hopper landmarks (red and orange). B) SimBA ROI positioning for training of hopper (red arrow) and cotton bud (blue arrow) aligned by food hopper bars shown by white arrows.

### Behavioural classifiers training using SimBA

To define the behaviour of interest, sniffing of introduced odour cues Simple Behavioural Analysis was used (SimBA v 2.0.7). A total of 80 videos were used for this step of the analysis pipeline, as per SimBA guidelines (Goodwin et al., 2024), this included the 60 videos used for DLC training. The videos were pre-processed for SimBA training by removing frames which did not contain the cotton bud (odour cue presentation), these were frames that had DLC pose estimation likelihood output for the cotton bud of <0.8.

Subsequently DLC pose estimation for the body parts was fed into SimBA for feature extraction, inputting only the mouse body parts identified from DLC (Figure 2A blue, green and purple dots), and SimBA annotated regions of interest (ROIs) (Figure 2B). This was combined with manual scoring of sniffing performed in Simple Video Coder, transformed to Solomon format using a custom-made python scripts and imported into SimBA (all data and scripts available on github <u>sboya23/ML-analysis-olfaction</u>). Within the SimBA workflow the DLC body pose estimation data was used without outlier correction, and manual scoring data was used to train SimBA to recognise two behavioural classifiers – “sniffs” and “does not sniff”. Removing frames without cotton bud presentation reduced the size of the dataset and improved the balance between “sniffs” and “does not sniff” annotation, improving the accuracy of classifications.

In SimBA, ROIs (metal hopper and the cotton bud) were applied to aid feature extraction in each frame (Figure 2B). For the training phase the ROIs were set in a single video, then applied to all videos in the training set, and the size of the ROIs was standardised according to the first video in which the ROIs were set. Features extraction, features extraction appended to body parts, and extra sub-feature extraction (within animal 3 body part angles, and frame by frame body part movement) were used to train a “sniffs” and a “does not sniff” behavioural classifier models. For list of features used see <u>ML-analysis-olfaction/SimBA-training at main ·</u> <u>sboya23/ML-analysis-olfaction</u>. The models were trained with the following settings – random forest algorithm with 2000 estimators, 80% train-20% test spit was used. The training was performed with two learning curve splits (see <u>ML-analysis-olfaction/SimBA-training at main ·</u> <u>sboya23/ML-analysis-olfaction</u>). The models showed precision = 0.92, recall = 0.86, and f1 score = 0.89 for “sniffs”, and precision = 0.98, recall = 0.99, and f1 score = 0.98 for “does not sniff”. Optimised detection threshold for “sniffs” and “does not sniff” was 0.5 confidence, and 200 ms duration of the behavioural bout.

### Validation of the model on unseen videos

To validate the devised pipeline, the trained DLC and SimBA models were used to determine sniffing times in a set of unseen test videos, that had been previously manually scored (Boyanova et al., 2025). The videos were first analysed using the trained DLC model for pose estimation. The body parts output was fed in a new SimBA project. SimBA ROIs were positioned for each video, the ROI size was standardised according to a randomly selected video, and features were extracted as during the training phase. The previously trained SimBA models classified both “sniffs” and “does not sniff” bouts (0.5 confidence threshold and 200 ms bout length), mutual exclusivity of “sniffs” and “does not sniff” was applied where “sniffs” was set as the winner with threshold 0 to ensure mutually exclusive classifications in favour of “sniffing”. The time spent sniffing was determined by the model in 1 s time bins, which were then subdivided for the nine odour presentations into time periods of interest (TOI) for each video, to generate the time spent sniffing of each odour cue (three odours presented three times for an equal amount of time adjusted to the individual length of each video).

### Statistical analysis

Pearson R correlation in R Studio (R version 4.4.2) was used to measure the correlation between manual and ML scoring of videos, and between two manual scorers. The generalised linear mixed effects models (glmer) and linear mixed-effects models (lmer) and from the lme4 and lmerTest packages (Bates, Mächler, Bolker, & Walker, 2015; Kuznetsova, Brockhoff, & Christensen, 2017) in R Studio (R version 4.4.2) were used for data analysis, based on (Colom-Cadena et al., 2023; ‘Spires-Jones Lab GitHub’, 2025). To assess if data obtained from the manual scoring compared with the ML method was filtered differently, a glmer model with binominal distribution: *status ∼ source*age with individual video as a random effect*, where status means whether a video would be filtered or not (0, 1), and source – whether the video was scored using manual or ML method was used. If the mouse spent less than 10 seconds interacting with the first presentation of a given odour, this run and the consecutive two presentations of the same odour were filtered out. We performed analysis of the effect of age using the average age of the mice at every time point; thus, we had two age groups for each study (*C9orf72* versus wildtype controls and *Tardbp* versus wildtype). Fixed factors and interactions were assessed using Anova Type III Wald Chi square tests. To assess whether there is a significant difference in the values obtained from manual or ML scoring, and to assess the effects of genotype, odour type presentation and sex we used the lmer model *time spent sniffing ∼ source*genotype*(odour type presentation + sex) with the individual video as a random effect* for both studies. Odour type presentation was defined as all three presentations of each odour – water, familiar mouse and novel mouse odour, and the two age groups were analysed separately for each mouse model (*C9orf72* versus wildtype controls and *Tardbp* versus wildtype). Factor significance was assessed using Type III ANOVA with Satterthwaite’s method and is reported (Colom-Cadena et al., 2023). For this task, analysis of performance can only be measured if sufficient engagement with the odour cues occurs at the first presentation, thus data must be filtered to remove experiments which do not meet this requirement. Diagnostics for glmer and lmer were performed by applying the DHARMa diagnostics package (Hartig, Lohse, & leite, 2024), and residuals plotting, respectively. The emmeans package with Bonferroni correction was used for post hoc analysis of comparisons of interest when a significant main effect of a variable was observed to identify which groups differed. Python and R script generation and refinement was assisted by the large language model ChatGPT.

## Results

### Correlation between ML and manual sniffing time in the olfactory habituation-dishabituation task

To determine the efficacy of our ML pipeline to quantify the time spent interacting with odour cue (sniffing) in the olfactory habituation-dishabituation task, we performed correlation analysis between manual scoring and the output of the ML analysis pipeline. We evaluated our pipeline using previously reported data (Boyanova et al., 2025) comparing colony-matched wildtype (WT) and *C9orf72^GR400/+^* mice, and colony-matched WT and *Tardbp^Q331K/Q331K^* mice that model different aspects of ALS/FTD, which were manually scored. These data are longitudinal and measured olfaction in young (15-weeks of age) and old (67-weeks of age) mice.

Comparing the time spent sniffing in WT and *C9orf72^GR400/+^* at 15- and 67-weeks of age, we observed strong and significant positive correlation between manual and ML pipeline determined sniffing time – Pearson r = 0.90, R^2^ = 0.82, p = 9.94e-95 (Figure 3A), and Pearson r = 0.87, R^2^ = 0.76, p = 6.79e-51 (Figure 3B), respectively. For the WT and *Tardbp^Q331K/Q331K^* mice we observed a similar correlation between manual and ML determined sniffing times, at both 15- and 67-weeks of age – Pearson r = 0.93, R^2^ = 0.86, p = 2.42e-121 (Figure 3C), and Pearson r = 0.88, R^2^ = 0.77, p = 8.81e-59 (Figure 3D). The correlation between two manual scorers is comparable (Figure 3E) to the correlation between ML and manual methods, supporting the efficacy of our ML pipeline for the quantification of sniffing time in the habituation-dishabituation task.

**Figure 3.**
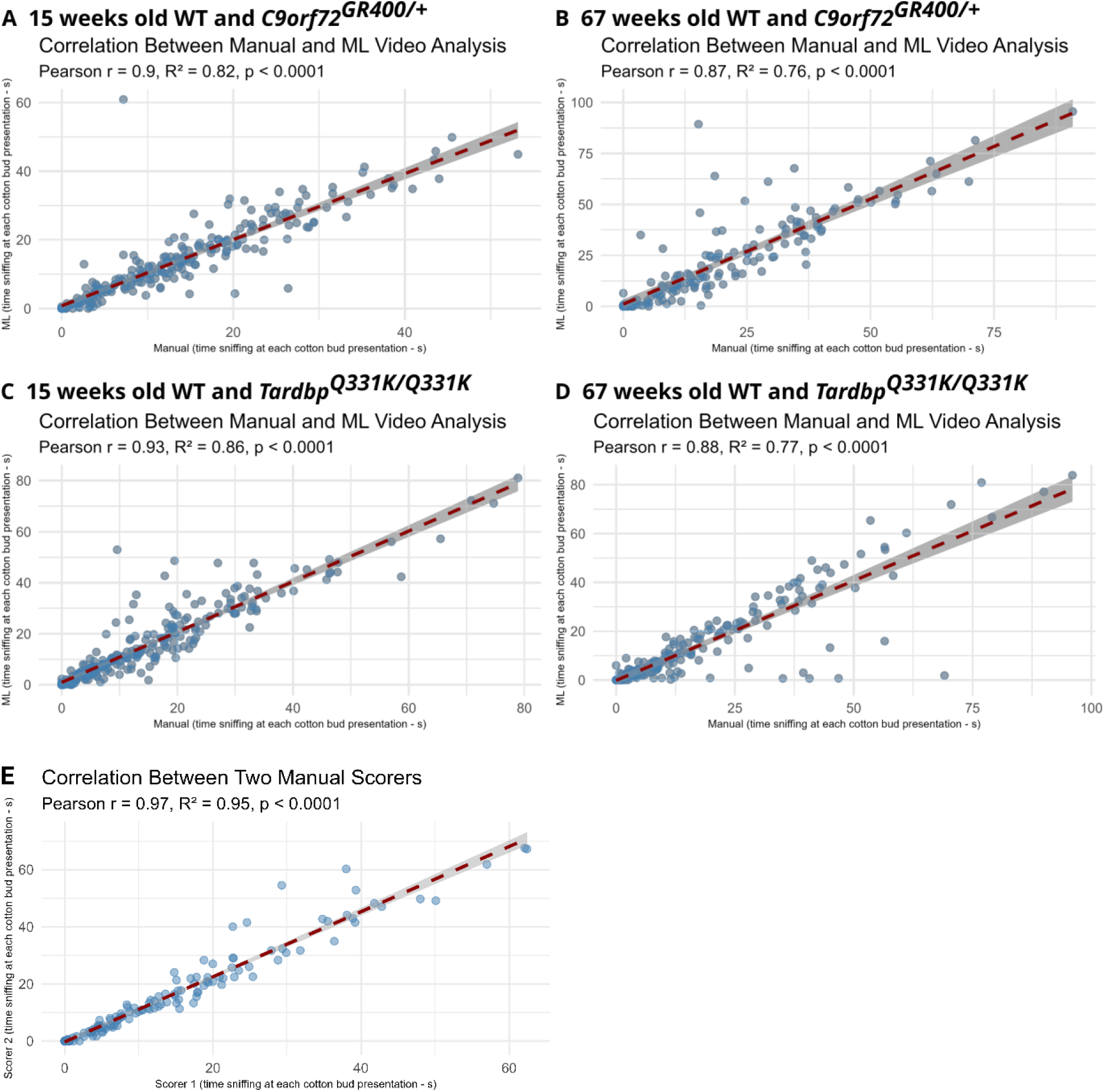
Relationship between sniffing annotation by automated pipeline and manual annotation. Correlation between manual and ML estimation of the time spent sniffing each odour presentation in A) 15-weeks of age (Pearson r = 0.90, R^2^ = 0.82, p = 9.94e-95) and B) 67-weeks of age (Pearson r = 0.87, R^2^ = 0.76, p = 6.79e-51) in WT and *C9orf72^GR400/+^* mice, and in C) 15-weeks of age (Pearson r = 0.93, R^2^ = 0.86, p = 2.42e-121), and D) 67-weeks of age (Pearson r = 0.88, R^2^ = 0.77, p = 8.81e-59) in WT and *Tardbp^Q331K/Q331K^* mice. For number of videos, see Supplementary Table 1. E) Correlation between two manual scorers of the time spent sniffing each odour presentation (Pearson r = 0.97, R^2^ = 0.95, p = 2.75e-92). Total of 16 videos were selected at random from the *C9orf72* and the *Tardbp* studies, four at each age (15- and 67-weeks of age).

### Comparison of ML and manual quantification for the assessment of technical and biological effects in the olfactory habituation-dishabituation task

To analyse the olfactory habituation-dishabituation dataset for the biological effects of sex, age and genotype of animals, trials in which animals did not engage with the task, as indicated by < 10 seconds sniffing of the first presentation of each odour, are excluded (‘Behavioral Models | Behavioral and Functional Neuroscience Lab | Wu Tsai Neurosciences Institute’, 2023). Therefore, we first determined if our ML pipeline created bias during this analysis step. We ran a fitted generalised linear mixed model – *status of filtering (unfiltered or filtered on first presentation of odour <10 seconds) ∼ source (manual or ML)*age with video ID as a random factor*. For the *C9orf72* study, we did not observe an effect of **source** of data (manual or ML) (χ^2^ = 0.0109, df = 1, p = 0.91675), **age** (χ^2^ = 1.2413, df = 1, p = 0.26522) or **source-age interaction** (χ^2^ = 1.7922, df = 1, p = 0.18066) on the relative filtering of the data (Figure 4A). However, for the *Tardbp study* we observed a **source of data (manual or ML) - age interaction** (χ^2^ = 5.2677, df = 1, p = 0.02173) on the filtering of the data. Post hoc analysis using Bonferroni correction showed a difference between filtering status for ML and manual data at the 67-week time point (p = 0.0021). No main effects of **age** (χ^2^ = 2.2131, df = 1, p = 0.13685) or **source of the data** (manual or ML) (χ^2^ = 0.4906, df = 1, p = 0.48368) on filtering were observed in the *Tardbp* study (Figure 4B).

**Figure 4:**
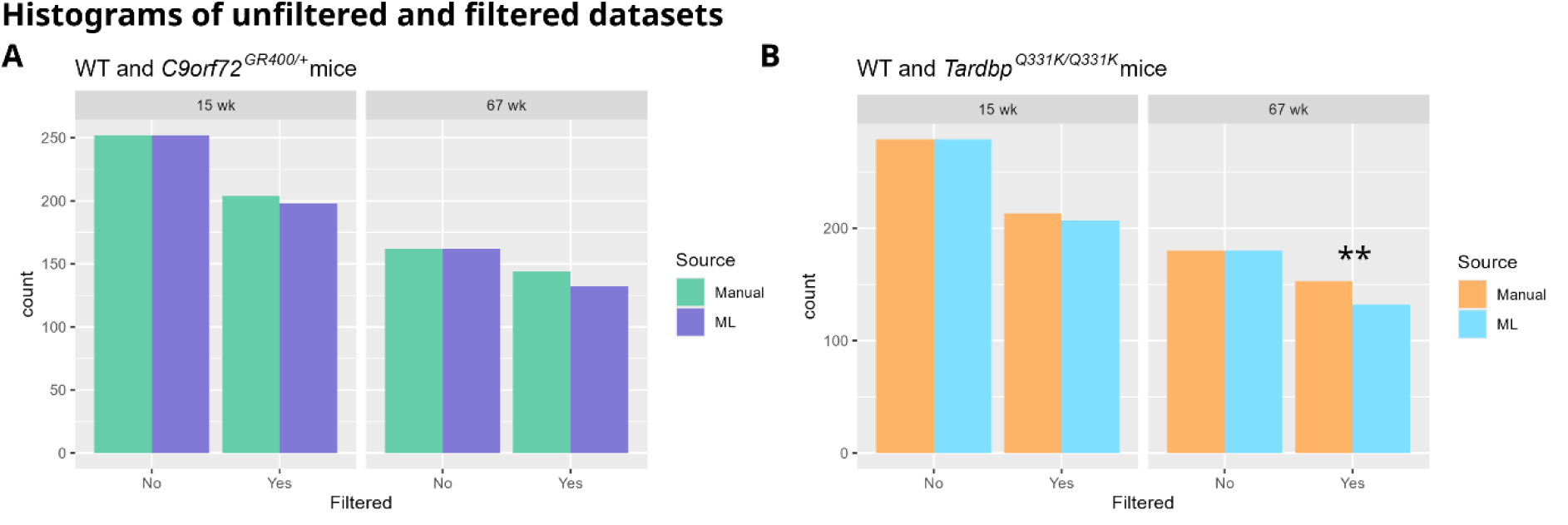
Comparison of data filtering from manual annotation and automated pipeline. Histograms showing the number of data points in the unfiltered and filtered (total sniffing time >10 seconds, first presentation of odour) datasets. A) *C9orf72^GR400/+^* study, green – manual dataset (n = 252 non-filtered, n = 204 filtered 15-weeks of age, n=162 non-filtered, n = 144 filtered 67-weeks of age), purple – ML dataset (n = 252 non-filtered, n = 198 filtered 15 weeks, n = 162 non-filtered, n = 132 filtered 67 weeks). B) *Tardbp^Q331K/Q331K^*study, orange – manual dataset (n = 279 non-filtered, n = 213 filtered 15-weeks of age, n = 180 non-filtered, n = 153 filtered 67 weeks), blue – ML dataset (n = 279 non-filtered, n = 207 filtered 15 weeks, n = 180 non-filtered, n = 132 filtered 67-weeks of age). Effect of source of data (manual or ML) and age interaction χ^2^ = 5.2677, df = 1, p = 0.02173, ** p<0.01 post hoc analysis with Bonferroni correction.

To determine if manual compared with ML annotation of sniffing behaviour altered technical or biological outcomes of these olfaction studies, we assessed the filtered data for the effect of data source (manual vs ML) on the time spent interacting with the odour cue across the task by including of data source in a lmer (*time spent sniffing ∼ source*genotype*( odour type presentation + sex) + (1|video ID)*). We also determine the effects of genotype, sex and odour type presentation order. In the case of significant main effects, we explored post hoc effects of interest between data sources (manual or ML) to compare the performance of the two pipelines. We found no evidence that data source affected the statistical outcome in either mouse model, at either time point. Moreover, data source did not interact with technical (odour (water, familiar, social) and presentation event (1-9)), or biological (sex, genotype) variables (for full statistical output see Supplementary Table 2) (Figure 5). Overall, our ML pipeline recapitulated parameters to control for task efficacy and to test our primary hypothesis that mouse genotype affected odour habituation-dishabituation; despite the effect of ML on filtering rate at the 67-week time point in the *Tardbp* study.

**Figure 5:**
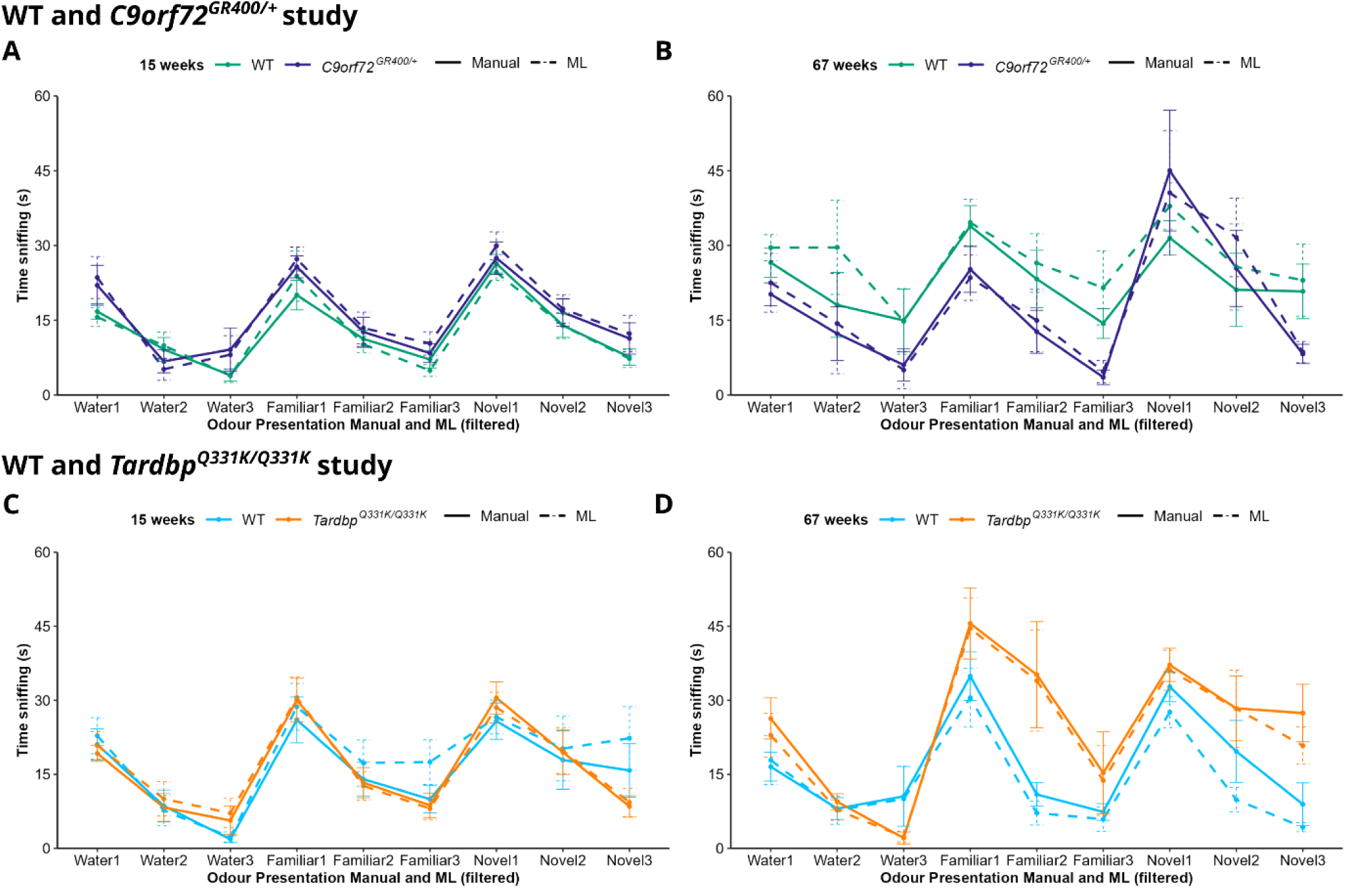
Comparison of the time spent sniffing determined by manual or ML methods by order of odour presentation and genotype. A-D) Manual scoring (solid line) or the machine learning pipeline (dashed line) generated data of time sniffing each presentation (1-3) of control (water), social familiar and social novel odours, in colony matched WT control (green) and *C9orf72^GR400/+^* (purple), and colony matched WT control (blue) and *Tardp^Q331K/Q331K^* mice (orange), at (A, C) 15 weeks and (B, D) 67-weeks of age. A) No effects of data source (manual or ML) F (1, 342.90) = 0.988, p = 0.75345, an effect of genotype F (1, 25) = 7.3660, p = 0.01187, sex F (1, 24.73) = 4.4478, p =0.04525, odour type presentation F (8,342.27) = 41.3998, p < 2e-16 was observed at 15-weeks of age in the *C9orf72^GR400/+^*study. B) No effect of data source (manual or ML) F (1, 222.863) = 0.9682, p = 0.32620, an effect of odour type presentation F (8, 223.135) = 14.5512, p < 2e-16, genotype*odour type presentation F (8, 223.135) = 2.3644, p = 0.01843 was observed at 67-weeks of age in the *C9orf72^GR400/+^* study. C) In WT and *Tardp^Q331K/Q331K^*mice at 15 weeks of age, no effect of data source (manual or ML) F (1, 356.02) = 2.6738, p = 0.1029, an effect of odour type presentation F (8, 359.25) = 28.2165, p < 2e-16, and a genotype* odour type presentation interaction F (8, 359.25) = 2.3737, p = 0.0168 were observed. D) In WT and *Tardp^Q331K/Q331K^* mice, at 67 weeks of age, no effect of data source (manual or ML) F (1,232.057) = 3.3788, p = 0.06732, an effect of odour type presentation F (8, 230.995) = 24.1319, p < 2.2e-16, significant genotype* odour type presentation interaction F (8, 230.995) = 4.8783, p = 1.41e-05 was observed. Error bars represent mean ± SEM – standard error mean. At 15 weeks WT (controls for *C9orf72^GR400/+^*) n = 14, Female n = 5; *C9orf72^GR400/+^* n = 14, Female n = 8; WT n = 16, Female n = 8 (controls for *Tardp^Q331K/Q331K^*). *Tardp^Q331K/Q331K^* n = 15, Female n = 6. At 67 weeks WT (controls for *C9orf72^GR400/+^*) n = 11, Female n = 5; *C9orf72^GR400/+^* n = 7, Female n = 3; WT n = 11, Female n = 7 (controls for *Tardp^Q331K/Q331K^*). *Tardp^Q331K/Q331K^* n = 9, Female n = 3.

To further assess the ML pipeline’s utility, we undertook post hoc comparisons for any significant main effects observed in the lmer. We particularly assessed odour type presentation order to determine if olfactory habituation and dishabituation was detected, a key technical parameter for the validity of the experiment. We undertook this for every genotype at each age, and source of data (ML or manual), to determine if ML-generated data resulted in the same technical and biological conclusions as manually annotated data that we previously reported.

For the *C9orf72* study at 15 weeks of age we detected a main effect of **genotype** (F (1, 25.00) = 7.3660, p = 0.01187), **odour type presentation** (F (8, 342.27) = 41.3998, p < 2e-16), and **sex** (F (1, 24.73) = 4.4478, p = 0.04525) (Figure 5A). The post-hoc comparisons of both the manual and the ML data detected significant habituation to social odours, and significant dishabituation from control to social odours as well as dishabituation between social odours in both the WT and *C9orf72^GR400/+^* mice. Additionally, manual annotation and the ML pipeline found that the *C9orf72^GR400/+^* mice also show significant habituation to the control odour, and disinhibition from familial social odour to control odour (Supplementary Table 2A). The only post hoc discrepancy between the manual data and our ML pipeline was that the manual scoring showed habituation of the WT mice to the control odour, whereas the ML pipeline did not detect this, at 15-weeks of age.

For the *C9orf72* study at 67-weeks of age, we observed a main effect of **odour type presentation order** F (8, 223.135) = 14.5512, p < 2e-16, and an interaction between **genotype and odour presentation** F (8, 223.135) = 2.3644, p = 0.01843 (Figure 5B). Post hoc comparisons of both manual annotation and the ML pipeline data failed to detect significant habituation to novel social odour and the control odour in the WT, and to familiar social odour and control in the *C9orf72^GR400/+^* mice but detected significant habituation to the novel social odour in the *C9orf72^GR400/+^* mice, at the later timepoint. Manual annotation and ML data output successfully detected dishabituation from control to novel social odour, and from familiar to novel social odour in the *C9orf72^GR400/+^* mice, and dishabituation from control to familiar social odour in WT mice. However, at 67-week-of age manual annotation detected significant habituation to the familiar social odour in the WT mice, but this was not detected using the ML pipeline. Whereas the ML pipeline detected dishabituation from control to novel social odour, which was not detected by manual scoring in WT mice (Supplementary Table 2B). Importantly, no evidence of genotype effect (by post-hoc comparison using Bonferroni correction for multiple comparisons) on either habituation or dishabituation was observed using either manually or ML-extracted data in the *C9orf72^GR400/+^* study. Thus, ML-extracted data produced the same overall biological conclusions as manually annotated data.

In the *Tardbp* study, at 15 weeks of age we detected a main effect of **odour type presentation order** F (8, 359.25) = 28.2165, p < 2e-16, and an interaction between **genotype and odour type presentation** F (8, 359.25) = 2.3737, p = 0.0168. Post hoc comparisons detected habituation to the control odour in WT mice, and to the familiar, and the novel social odour in the *Tardbp^Q331K/Q331K^*mice, using data extracted by both manual annotation and the ML pipeline. Similarly, both data extraction methods detected dishabituation from control to the social smells in the WT and *Tardbp^Q331K/Q331K^* mice, and dishabituation between the social smells in the *Tardbp^Q331K/Q331K^* mice. The only discrepancies between data source (manual or ML) were observed in WT mice, at the younger time point; in which manual annotation detected habituation to familiar social smell, and dishabituation from familiar to novel smell, whereas the ML pipeline did not (Supplementary Table 2C). No evidence of genotype effect (by post-hoc comparison) on either habituation or dishabituation was observed using either manually extracted or ML-generated data in the *Tardbp* study at the younger timepoint.

In the *Tardbp* study, at 67-weeks of age, we detected an effect of **odour type presentation** F (8, 230.995) = 24.1319, p < 2.2e-16, and an interaction between **genotype and odour type presentation** F (8, 230.995) = 4.8783, p = 1.41e-05. At 67-weeks of age, habituation to familiar odour in the WT, and *Tardbp^Q331K/Q331K^* mice, and habituation to novel social odour in the WT was detected using our ML pipeline, consistent with previous manual annotation. Similarly, dishabituation from familiar to novel social odour in the WT and the *Tardbp^Q331K/Q331K^*mice was detected by ML as previously reported using manual annotation. Also, dishabituation from control to social odours in the *Tardbp^Q331K/Q331K^* mice was detected using both methods of data generation. However, the previously detected, using manual annotation, dishabituation from novel to familiar social odour, and from control to social odours in the WT was not detected in our ML pipeline. Our ML method was also not able to detect habituation to water in the *Tardbp^Q331K/Q331K^* mice but did detect dishabituation between social odours which was previously not detected using manual analysis. When we explored post hoc comparison between genotypes, both data extraction methods (manual and ML) showed significant genotype difference only at the second presentation of control odour (manual p = 0.0277, ML p = 0.0026), at the later time-point in the *Tardbp^Q331K/Q331K^* study.

Overall, data extracted using our ML pipeline recapitulated the key parameter to control for task efficacy (odour presentation order) and our primary biological hypothesis to test the effect of an animal’s genotype on odour habituation or dishabituation in the task. Moreover, our analysis found no significant effect of the source of data (manual or ML), supporting that data extracted using our new ML pipeline is comparable to data extracted using manual annotation.

We found some evidence of filtering differences and inconsistencies in the post-hoc analysis between the data sources, but these did not impact our overall assessment of the technical validation of the task or key biological outcomes in the two mouse models.

## Discussion

Here, we developed and validated a new ML pipeline to efficiently extract data from the mouse olfactory habituation-dishabituation task, using single camera side-view videos. We show that freely accessible and widely used ML tools can reach high and significant correlation to manual scoring even in complex home cage environments filmed from the side using a single camera, and extract data that leads to similar conclusions to manual annotation methods. This type of video orientation is compatible with long-term housing of animals under high-welfare (including group housing) IVC conditions, facilitating the acquisition of longitudinal behavioural data from animals held in the home environment. Moreover, the use of a single camera reduces data complexity compared with multiple camera views and circumvents the need for complex, computationally intensive data integration steps. Indeed, the ML pipeline reported here can be effectively run on a single desktop computer (Nvidia RTX4090 GPU), making it accessible to researchers who do not have access to high-performance clusters and reducing the energy impact of this type of automated analysis.

A number of considerations are required for implementation of this ML pipeline. Technical consistency, particular the placement and angle of the camera and task rig, and the orientation and length of the cotton bud odour cue during data acquisition are essential for the successful implementation of ML data extraction pipelines. Prior to the start of analysis, both these technical aspects and video data quality control should be undertaken, including the removal of any corrupted or interrupted videos or experiments in which technical errors occurred. Application of the pipeline to new datasets, requires a validation step using a subset of manually scored data (ideally by two independent scorers) to determine the efficacy of the pipeline to the new use case, and if retraining of either DLC or SimBA will be required. In particular, attention to the manual annotation of ROIs in SimBA for new datasets is required, to ensure consistency between users of the pipeline.

This ML pipeline has number of limitations, firstly the ROI annotated in SimBA applies a two-dimensional projection on three-dimensional video data, although this is adequate for the task automated in this study, this may confound the efficacy of this tool for other use cases. Our ML pipeline underscored sniffing behaviour in the *Tardbp* 67-week timepoint, although this issue did not affect the biological conclusion of the research. Thus, our ML pipeline does not extract behavioural task performance data identically under all conditions and revalidation is required prior to implementation in novel datasets. The development of this ML pipeline was enabled by access to two large longitudinal datasets, collected under identical technical conditions, which provided a large pool of high-quality training and test video data. Effective use of pose-estimation and SimBA for more complex use cases (as reported here) requires adequately powered datasets, for train-test cycles and validation steps, which may not be available to all researchers. In particular, the lack of standardisation of data acquisition methods between research groups, and/or small studies may limit the broader utility of these methods. Use of alternative methods of animal tracking such as segmentation, rather than pose estimation may reduce the size of training datasets, and partially address this limitation and would be an important avenue for future investigation (Chan et al., 2025).

Further future improvements for this olfactory habituation-dishabituation ML pipeline would include developing classifiers which better differentiate between types of interaction with the odour cue (sniffing compared with licking or biting). Similarly, these tools can be further developed to classify and quantitate other behaviours in the home-cage environment, observed using a single-side-view infrared camera, enabling additional studies of innate and task-mediated mouse behaviour. In summary, here we demonstrate that effective and efficient ML pipelines can be generated to accurately and effectively automate the quantification of complex behaviours in mice using a single side-view projection with body part occlusion, enhancing research productivity.

## Acknowledgement

We thank Loukia Katsouri (Sainsbury Wellcome Centre), Marius Bauza (Sainsbury Wellcome Centre) and Julija Krupic (UK Dementia Research Institute-UCL) for their helpful input into the development of this pipeline. We thank the MRC Mary Lyon Centre husbandry, genotyping and phenotyping teams for their technical assistance in collecting data analysed in this study.

## Data availability

Primary data outputs, genotyping codes, python, and R scripts are available at GitHub: <u>sboya23/ML-analysis-olfaction</u>. Other relevant data are included in the article and its supplementary information. Raw data are available from F.K.W. upon reasonable request.

## Ethics approval statement for work involving animals

All animal studies were licensed by the Home Office under the Animals (Scientific Procedures) Act 1986 Amendment Regulations 2012 (SI 4 2012/3039), UK, and additionally approved by the Institutional Ethical Review Committee.

## Supplementary Materials

**Supplementary Table 1:** Number of mice used for the analysis in each test at each time point. Mice were excluded from analysis *only* if they had to be culled for welfare reasons unless otherwise specified. 2 videos were excluded from any stage of the training and analysis pipeline due to wrong hopper used (67 weeks - C9ORF72-GR400-MAT-B6J/1.3c, C9ORF72-GR400-MAT-B6J/2.2d)

|  |  |  |  |  |
| --- | --- | --- | --- | --- |
| WT and *C9orf72*<sup>GR400/+</sup><br>Olfaction test | 15 weeks | DLC Train (15 videos)<br>WT = 7 (4 female)<br>*C9orf72*<sup>GR400/+</sup> = 8 (3 female)<br>SimBA train (15 DLC videos + 5 additional videos)<br>WT = 10 (7 female)<br>*C9orf72*<sup>GR400/+</sup> = 10 (4 female) | 67 weeks<br>2 videos excluded due to procedural failure; wrong hopper used (C9ORF72-GR400-MAT-B6J/1.3c, C9ORF72-GR400-MAT-B6J/2.2d) | DLC Train (15 videos)<br>WT = 8 (6 female)<br>*C9orf72*<sup>GR400/+</sup> = 7 (4 female)<br>SimBA train (15 DLC videos + 5 additional videos)<br>WT = 9 (6 female)<br>*C9orf72*<sup>GR400/+</sup> = 11 (6 female) |
|  |  | Test (28 videos)<br>WT = 14 (5 female)<br>*C9orf72*<sup>GR400/+</sup> = 14 (8 female) |  | Test (18 videos)<br>WT = 11 (5 female)<br>*C9orf72*<sup>GR400/+</sup> = 7 (3 female) |
| WT and *Tardbp*<sup>Q331K/Q331K</sup><br>Olfaction test | 15 weeks<br>2 videos which were used for DLC training were excluded from classifiers training – acquisition technical error (wrong order of | DLC Train (15 videos)<br>WT = 8 (4 female)<br>*Tardbp*<sup>Q331K/Q331K</sup> = 7 (5 female)<br>SimBA train (20 videos)<br>WT = 11 (4 female) | 67 weeks<br>1 mouse excluded from the test phase analysis due to corrupted video file (TDP43-Q331K-B6J-IC/4.2e) | DLC Train (15 videos)<br>WT = 7 (3 female)<br>*Tardbp*<sup>Q331K/Q331K</sup> = 8 (5 female)<br>SimBA train (20 videos)<br>WT = 10 (4 female) |
|  | odours presented during the test) these were valid for training of body pose estimation. (TDP43-Q331K-B6J-IC/1.1c, TDP43-Q331K-B6J-IC/6.1d),<br>Mouse TDP43-Q331K-B6J-IC/4.1a was excluded from the final analysis because of a technical error during data acquisition (wrong order of odours presented during the test). | ***Tardbp*<sup>Q331K/Q331K</sup>**<br>**= 9 (3 female)** |  | ***Tardbp*<sup>Q331K/Q331K</sup>**<br>**= 10 (7 female)** |
|  |  | **Test (31 videos)**<br>**WT = 16 (8 females)**<br>***Tardbp*<sup>Q331K/Q331K</sup>**<br>**= 15 (6 female)** |  | **Test (20 videos)**<br>**WT = 11 (7 female)**<br>***Tardbp*<sup>Q331K/Q331K</sup>**<br>**= 9 (3 female)** |

**Supplementary Table 2:** Full statistical analysis output. The blue cells represent comparisons relevant to odour habituation, and the purple cells - to odour dishabituation, light and dark for manual and ML, respectively, grey signifies not significant post hoc comparison.

| **A: WT vs *C9orf72*<sup>GR400/+</sup> 15 weeks** |  |  |  |
| --- | --- | --- | --- |
| Source – not significant – F (1, 342.90) = 0.988, p = 0.75345 |  |  |  |
| Genotype F (1, 25) = 7.3660, p = 0.01187 |  |  |  |
| Odour type presentation F (8, 342.27) = 41.3998, p <2e-16 |  |  |  |
| Sex F (1, 24.73) = 4.4478, p = 0.04525 |  |  |  |
| Bonferroni correction: pairwise ~ odour type presentation \| genotype \| source |  |  |  |
| Mouse group | Odour and presentation comparison | P value (manual) | P value (ML) |
| WT | familiar odour presentation 1 (F1) vs familiar odour presentation 3 (F3) | 0.0137 | <.0001 |
|  | novel odour presentation 1 (N1) vs novel odour presentation 3 (N3) | <.0001 | <.0001 |
|  | water presentation 1 (W1) vs water presentation 3 (W3) | 0.0157 | 0.0757 |
|  | W3 – F1 | 0.0006 | <.0001 |
|  | W3 – N1 | <.0001 | <.0001 |
|  | F3 – N1 | <.0001 | <.0001 |
|  | F3 – W1 | 0.2403 | 0.1570 |
|  | N3 – F1 | 0.0293 | 0.0002 |
|  | N3 – W1 | 0.5041 | 1.0000 |
| *C9orf72*<sup>GR400/+</sup> | F1 – F3 | <.0001 | <.0001 |
|  | N1 – N3 | 0.0001 | <.0001 |
|  | W1 – W3 | 0.0267 | 0.0020 |
|  | W3 – F1 | <.0001 | <.0001 |
|  | W3 – N1 | <.0001 | <.0001 |
|  | F3 – N1 | <.0001 | <.0001 |
|  | F3 – W1 | 0.0185 | 0.0144 |
|  | N3 – F1 | 0.0002 | 0.0002 |
|  | N3 – W1 | 0.1808 | 0.0931 |

| Bonferroni correction<br>genotype * sex \| source | Manual scoring | ML scoring |
| --- | --- | --- |
| Females – contrast for genotype | 0.4374 | 0.0782 |
| Males contrast for genotype | 0.7915 | 0.7339 |
| WT contrast for sex | 0.4301 | 0.3038 |
| *C9orf72*<sup>GR400/+</sup> contrast for sex | 0.9461 | 1.0000 |

| **B: WT vs *C9orf72*<sup>GR400/+</sup> 67 weeks** |  |  |  |
| --- | --- | --- | --- |
| Source – not significant – F (1, 222.863) = 0.9682, p = 0.32620 |  |  |  |
| Odour type presentation – F (8, 223.135) = 14.5512, p < 2e-16 |  |  |  |
| Genotype * Odour type presentation – F (8, 223.135) = 2.3644, p = 0.01843 |  |  |  |
| Bonferroni correction: pairwise ~ odour type presentation \| genotype \| source |  |  |  |
| Mouse group | Odour and presentation comparison | P value (manual) | P value (ML) |
| WT | familiar odour presentation 1 (F1) vs familiar odour presentation 3 (F3) | 0.0190 | 0.6922 |
|  | novel odour presentation 1 (N1) vs novel odour presentation 3 (N3) | 1.0000 | 0.4085 |
|  | water presentation 1 (W1) vs water presentation 3 (W3) | 1.0000 | 0.8635 |
|  | W3 – F1 | 0.0175 | 0.0047 |
|  | W3 – N1 | 0.0701 | 0.0010 |
|  | F3 – N1 | 0.0816 | 0.1942 |
|  | F3 – W1 | 1.0000 | 1.0000 |
|  | N3 – F1 | 0.6863 | 1.0000 |
|  | N3 – W1 | 1.0000 | 1.0000 |
| *C9orf72*<sup>GR400/+</sup> | F1 – F3 | 0.3305 | 0.8283 |
|  | N1 – N3 | 0.0005 | 0.0011 |
|  | W1 – W3 | 1 | 1.0000 |
|  | W3 – F1 | 0.3908 | 0.5313 |
|  | W3 – N1 | <.0001 | 0.0006 |
|  | F3 – N1 | 0.0001 | 0.0009 |
|  | F3 – W1 | 1.0000 | 1.0000 |
|  | N3 – F1 | 1.0000 | 1.0000 |
|  | N3 – W1 | 1.0000 | 1.0000 |

| Bonferroni correction genotype * odour type presentation \| source<br>Genotype contrast at every odour and source | Manual scoring | ML scoring |
| --- | --- | --- |
| Water1 | 1.000 | 1.000 |
| Water2 | 1.000 | 1.000 |
| Water3 | 1.000 | 0.8017 |
| Familiar1 | 1.000 | 1.000 |
| Familiar2 | 1.000 | 1.000 |
| Familiar3 | 1.000 | 1.000 |
| Novel1 | 1.000 | 1.000 |
| Novel2 | 1.000 | 1.000 |
| Novel3 | 1.000 | 0.6330 |

| **C: WT vs *Tardbp*<sup>Q331K/Q331K</sup> 15 weeks** |  |  |  |
| --- | --- | --- | --- |
| Source – not significant – F (1, 356.02) = 2.6738, p = 0.1029 |  |  |  |
| Odour type presentation – F (8, 359.25) = 28.2165, p < 2e-16 |  |  |  |
| Genotype*Odour type presentation – F (8, 359.25) = 2.3737, p = 0.0168 |  |  |  |
| Bonferroni correction: pairwise ~ odour type presentation \| genotype \| source |  |  |  |
| Mouse group | Odour and presentation comparison | P value (manual) | P value (ML) |
| WT | familiar odour presentation 1 (F1) vs familiar odour presentation 3 (F3) | 0.0034 | 0.3029 |
|  | novel odour presentation 1 (N1) vs novel odour presentation 3 (N3) | 0.6751 | 1.0000 |
|  | water presentation 1 (W1) vs water presentation 3 (W3) | 0.0029 | 0.0004 |
|  | W3 – F1 | <.0001 | <.0001 |
|  | W3 – N1 | <.0001 | <.0001 |
|  | F3 – N1 | 0.0094 | 1.0000 |
|  | F3 – W1 | 1.0000 | 1.0000 |
|  | N3 – F1 | 0.4226 | 1.0000 |
|  | N3 – W1 | 1.0000 | 1.0000 |
| *Tardbp*<sup>Q331K/Q331K</sup> | F1 – F3 | 0.0001 | <.0001 |
|  | N1 – N3 | 0.0001 | 0.0013 |
|  | W1 – W3 | 0.1193 | 0.1636 |
|  | W3 – F1 | <.0001 | <.0001 |
|  | W3 – N1 | <.0001 | 0.0002 |
|  | F3 – N1 | <.0001 | 0.0002 |
|  | F3 – W1 | 0.8627 | 0.2183 |
|  | N3 – F1 | 0.0002 | 0.0002 |
|  | N3 – W1 | 1.0000 | 0.6959 |

| Bonferroni correction genotype * odour type presentation \| source<br>Genotype contrast at every odour and source | Manual scoring | ML scoring |
| --- | --- | --- |
| Water1 | 1.000 | 1.000 |
| Water2 | 1.000 | 1.000 |
| Water3 | 1.000 | 1.000 |
| Familiar1 | 1.000 | 1.000 |
| Familiar2 | 1.000 | 1.000 |
| Familiar3 | 1.000 | 0.6736 |
| Novel1 | 1.000 | 1.000 |
| Novel2 | 1.000 | 1.000 |
| Novel3 | 1.000 | 0.2590 |

| D: WT vs *Tardbp*<sup>Q331K/Q331K</sup> 67 weeks |  |  |  |
| --- | --- | --- | --- |
| Source – not significant – F (1,232.057) = 3.3788, p = 0.06732 |  |  |  |
| Odour type presentation – F (8, 230.995) = 24.1319, p < 2.2e-16 |  |  |  |
| Genotype*Odour type presentation – F (8, 230.995) = 4.8783, p = 1.41e-05 |  |  |  |
| Bonferroni correction: pairwise ~ odour type presentation \| genotype \| source |  |  |  |
| Mouse group | Odour and presentation comparison | P value (manual) | P value (ML) |
| WT | familiar odour presentation 1 (F1) vs familiar odour presentation 3 (F3) | 0.0012 | 0.0168 |
|  | novel odour presentation 1 (N1)<br>vs novel odour presentation 3 (N3) | 0.0020 | 0.0322 |
|  | water presentation 1 (W1) vs<br>water presentation 3 (W3) | 1.0000 | 1.0000 |
|  | W3 – F1 | 0.0071 | 0.1796 |
|  | W3 – N1 | 0.0092 | 0.1551 |
|  | F3 – N1 | 0.0014 | 0.0155 |
|  | F3 – W1 | 1.0000 | 1.0000 |
|  | N3 – F1 | 0.0023 | 0.0513 |
|  | N3 – W1 | 1.0000 | 1.0000 |
| *Tardbp*<sup>Q331K/Q331K</sup> | F1 – F3 | 0.0001 | <.0001 |
|  | N1 – N3 | 1.0000 | 0.6950 |
|  | W1 – W3 | 0.0203 | 0.1184 |
|  | W3 – F1 | <.0001 | <.0001 |
|  | W3 – N1 | <.0001 | <.0001 |
|  | F3 – N1 | 0.0340 | 0.0215 |
|  | F3 – W1 | 1.0000 | 1.0000 |
|  | N3 – F1 | 0.1120 | 0.0064 |
|  | N3 – W1 | 1.0000 | 1.0000 |

| Bonferroni correction genotype * odour<br>type presentation \| source<br>Genotype contrast at every odour and<br>source | Manual scoring | ML scoring |
| --- | --- | --- |
| Water1 | 1.0000 | 0.3506 |
| Water2 | 0.0277 | 0.0026 |
| Water3 | 1.0000 | 1.0000 |
| Familiar1 | 1.0000 | 1.0000 |
| Familiar2 | 1.0000 | 0.3316 |
| Familiar3 | 0.3195 | 0.5694 |
| Novel1 | 1.0000 | 1.0000 |
| Novel2 | 1.0000 | 1.0000 |
| Novel3 | 0.8011 | 1.0000 |

## References

Bates, D., Mächler, M., Bolker, B., & Walker, S. (2015). Fitting Linear Mixed-Effects Models Using lme4. Journal of Statistical Software, 67(1). 10.18637/jss.v067.i01

Behavioral Models | Behavioral and Functional Neuroscience Lab | Wu Tsai Neurosciences Institute. (2023, October 19). Retrieved 27 November 2025, from https://neuroscience.stanford.edu/shared-resources/behavioral-functional-lab/services-and-rates/behavioral-models

Boyanova, S., Banks, G., Lipina, T. V., Bains, R. S., Forrest, H., Stewart, M., … Wiseman, F. K. (2025). Multi-modal comparative phenotyping of knock-in mouse models of frontotemporal dementia/amyotrophic lateral sclerosis. Disease Models & Mechanisms, 18(8), dmm052324. 10.1242/dmm.052324

Chan, A. H. H., Putra, P., Schupp, H., Köchling, J., Straßheim, J., Renner, B., … Kano, F. (2025). YOLO-Behaviour: A simple, flexible framework to automatically quantify animal behaviours from videos. Methods in Ecology and Evolution, 16(4), 760–774. 10.1111/2041-210X.14502

Colom-Cadena, M., Davies, C., Sirisi, S., Lee, J.-E., Simzer, E. M., Tzioras, M., … Spires-Jones, T. L. (2023). Synaptic oligomeric tau in Alzheimer’s disease—A potential culprit in the spread of tau pathology through the brain. Neuron, 111(14), 2170–2183.e6. 10.1016/j.neuron.2023.04.020

Goodwin, N. L., Choong, J. J., Hwang, S., Pitts, K., Bloom, L., Islam, A., … Golden, S. A. (2024). Simple Behavioral Analysis (SimBA) as a platform for explainable machine learning in behavioral neuroscience. Nature Neuroscience, 27(7), 1411–1424. 10.1038/s41593-024-01649-9

Hartig, F., Lohse, L., & de leite, M. S. (2024). DHARMa: Residual Diagnostics for Hierarchical (Multi-Level / Mixed) Regression Models. Retrieved from https://cran.r-project.org/web/packages/DHARMa/index.html

Ison, J. R., Allen, P. D., & O’Neill, W. E. (2007). Age-related hearing loss in C57BL/6J mice has both frequency-specific and non-frequency-specific components that produce a hyperacusis-like exaggeration of the acoustic startle reflex. Journal of the Association for Research in Otolaryngology: JARO, 8(4), 539–550. 10.1007/s10162-007-0098-3

Kuznetsova, A., Brockhoff, P. B., & Christensen, R. H. B. (2017). lmerTest Package: Tests in Linear Mixed Effects Models. Journal of Statistical Software, 82(13). 10.18637/jss.v082.i13

Marks, M., Qiuhan, J., Sturman, O., von Ziegler, L., Kollmorgen, S., von der Behrens, W., … Yanik, M. F. (2022). Deep-learning based identification, tracking, pose estimation, and behavior classification of interacting primates and mice in complex environments. Nature Machine Intelligence, 4(4), 331–340. 10.1038/s42256-022-00477-5

Mathis, A., Mamidanna, P., Cury, K. M., Abe, T., Murthy, V. N., Mathis, M. W., & Bethge, M. (2018). DeepLabCut: Markerless pose estimation of user-defined body parts with deep learning. Nature Neuroscience, 21(9), 1281–1289. 10.1038/s41593-018-0209-y

Milioto, C., Carcolé, M., Giblin, A., Coneys, R., Attrebi, O., Ahmed, M., … Isaacs, A. M. (2024). PolyGR and polyPR knock-in mice reveal a conserved neuroprotective extracellular matrix signature in C9orf72 ALS/FTD neurons. Nature Neuroscience, 27(4), 643–655. 10.1038/s41593-024-01589-4

Pereira, T. D., Tabris, N., Matsliah, A., Turner, D. M., Li, J., Ravindranath, S., … Murthy, M. (2022). SLEAP: A deep learning system for multi-animal pose tracking. Nature Methods, 19(4), 486–495. 10.1038/s41592-022-01426-1

Rodgers, S. P., Born, H. A., Das, P., & Jankowsky, J. L. (2012). Transgenic APP expression during postnatal development causes persistent locomotor hyperactivity in the adult. Molecular Neurodegeneration, 7(1), 28. 10.1186/1750-1326-7-28

Ryan, B. C., Young, N. B., Moy, S. S., & Crawley, J. N. (2008). Olfactory cues are sufficient to elicit social approach behaviors but not social transmission of food preference in C57BL/6J mice. Behavioural Brain Research, 193(2), 235–242. 10.1016/j.bbr.2008.06.002

Spires-Jones Lab. (2025, November 28). Retrieved 28 November 2025, from GitHub website: https://github.com/Spires-Jones-Lab

Von Ziegler, L., Sturman, O., & Bohacek, J. (2021). Big behavior: Challenges and opportunities in a new era of deep behavior profiling. Neuropsychopharmacology, 46(1), 33–44. 10.1038/s41386-020-0751-7

White, M. A., Kim, E., Duffy, A., Adalbert, R., Phillips, B. U., Peters, O. M., … Sreedharan, J. (2018). TDP-43 gains function due to perturbed autoregulation in a Tardbp knock-in mouse model of ALS-FTD. Nature Neuroscience, 21(4), 552–563. 10.1038/s41593-018-0113-5

